# Scaling laws of molecular residence time

**DOI:** 10.1101/2024.02.05.578884

**Authors:** Shiyi Qin, Zhi Yang, Kai Huang

**Author notes:** Correspondence (K.H.). These authors contribute equally (Shiyi Qin, Zhi Yang).

## Abstract

Understanding the molecular residence autocorrelation function in liquid is of fundamental importance in physical and life science. Encoded in this function is not only the binding properties, but also the information of the liquid environment. Based on extensive in silico experiments and theoretical analysis, we reveal that power law residence scaling arises in both passive and active liquid, in contrast to the common sense of exponential decay. In simple homogeneous liquid, the scaling exponent depends solely on the system dimensionality. Such scaling law is robust against the superposition of diverse binding energies in single-phase liquid but can be breached if the system undergoes phase separation. Remarkably, in a dissipative system where phase separation is subject to non-equilibrium feedback controls, an anomalous power law emerges whose scaling exponent is in line with the puzzling residence scaling of transcription factors reported in recent experiments. Our results highlight the sensitivity of molecular residence to its surrounding liquid and suggest that active phase separation can serve as a scaling proofreading mechanism in gene regulation.

## Introduction

Unlike particles in solid that have fixed neighbors, liquid molecules regularly update their neighboring lists as they engage in transient interactions driven by noncovalent forces, such as electrostatic attraction, hydrogen bonding, van der Waals forces, pi-pi interactions, and the hydrophobic effect. Consider two liquid molecules that happen to reside near each other at a given moment. The likelihood of finding them in proximity at later moment decays with time, as described by a residence time autocorrelation function (RAF):

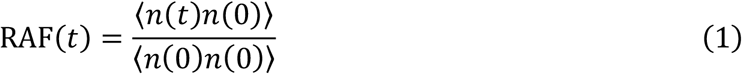

where *n*(*t*) is the number of proximal molecules at moment *t*. Understanding and characterizing this function is not only a classic problem in chemical physics, but also holds paramount importance in unraveling the inner workings of biological systems. For example, residence of transcription factors (TFs) at cis-regulatory elements (CREs) is needed to activate gene transcription^1–10^. Therefore, how long TFs can stay near their cognate sites is a fundamental question in biophysics.

The advancements of experimental technologies have yielded unprecedented insights into the RAF of TFs^7,11^. Nevertheless, the interpretation of the collected data depends on the choice of mathematical models and underlying physical assumptions^7^. It is widely believed in biophysics that, given a certain binding energy, the RAF should decay exponentially. This common sense has not been questioned even when a power law of RAF that strongly deviates from exponential decay was recently discovered by TF experiments^7,12–16^. Instead, a continuum of affinities model has been developed to explain the observed power law of RAF by the superposition of exponential RAFs^7,15,16^. However, the tenet of exponential RAF may fall short in capturing the complexity arising from the interplay between the binding kinetics and the molecular dynamics in liquid.

When the liquid has biological activities, one needs to further consider the nonequilibrium effects of biochemical reactions on the molecular dynamics. Yet another layer of complexity comes from the fact that biochemical reactions are not happening homogeneously in space. It has been now widely recognized that the cellular space is compartmentalized into various membraneless organelles (MLOs)^17–21^ via the process of liquid-liquid phase separation (LLPS)^22–30^. How the multi-phase liquid environment in biological systems could affect the RAF of biomolecules stands as a pressing question.

Here we design in silico experiments and apply scaling theory to systematically study the RAFs in both simple and complex liquid, in search of emergent scaling laws. Computer simulation has been a powerful tool to investigate molecular autocorrelation functions^31–33^. However, accurately calculating the RAF is computationally expensive. Moreover, the periodic boundary condition of the simulation box can lead to artifact of RAF at long-time scale (Fig. S2). We address these problems and accelerate the data processing by about three orders of magnitudes via optimizing the molecular neighboring analysis (see the Methods for more details). By using this technique, we obtain high-quality statistics of the RAFs across multiple timescales. Our results unambiguously show that power law is the rule rather than exception for long-time behavior of RAF of liquid molecules. In simple liquid, system dimension governs the scaling exponent of the power law. Diversifying the molecular binding energy and other perturbations of the system such as adding crowders can only change the amplitude of the power law but not the scaling exponent, as long as the liquid stays homogenous. We then investigate the effects of molecular aggregation and phase separation on the RAF and find that the latter can lead to strong deviations from the power law. Next, we construct and study an active droplet to explain the anomalous scaling law of TFs observed in experiments. We end the paper by discussing the biological function of this emergent scaling behavior in dissipative system.

### Universal scaling law in simple liquid

We start with all-atom molecular dynamics (MD) simulations of associated liquid rich in hydrogen bonds, such as water and alcohol. The RAFs of hydrogen bonding in systems of different dimensions are analyzed and shown in Fig. 1A-1C. In these log-log plots, all the RAFs are clearly non-exponential with long-time tails following power law functions. The scaling exponents of the power law functions unequivocally align with − *D*/2, where *D* is the dimension of the system. The dimension-dependent power law scaling is universal, as it does not depend on the type of the associated liquid or the specific definition of its hydrogen bonding (Fig. S1). The observed universality of the RAF scaling could be attributed to the conservation laws such as mass conservation (governing diffusion) and momentum conservation (governing hydrodynamics), both of which are implemented in all-atom MD. Using Brownian dynamics (BD) simulations without hydrodynamics, we further show that diffusion alone can preserve the dimension-dependent scaling law of RAF (Fig. 1D-1F). The high efficiency of the BD simulation allows us to systematically study the dependence of RAF on a wealth of factors including viscosity (Fig. 2A), crowding (Fig. 2B), molecular weight (Fig. 2C), binding strength (Fig. S4), and residence range (Fig. S5). Our results reveal that all these factors can amplify the power law of RAF. Therefore, in biological liquid where many of the above factors are simultaneously satisfied, non-exponential RAF with significant long-time tail is expected to be an intrinsic property of biomacromolecules such as the TFs.

**Fig. 1.**
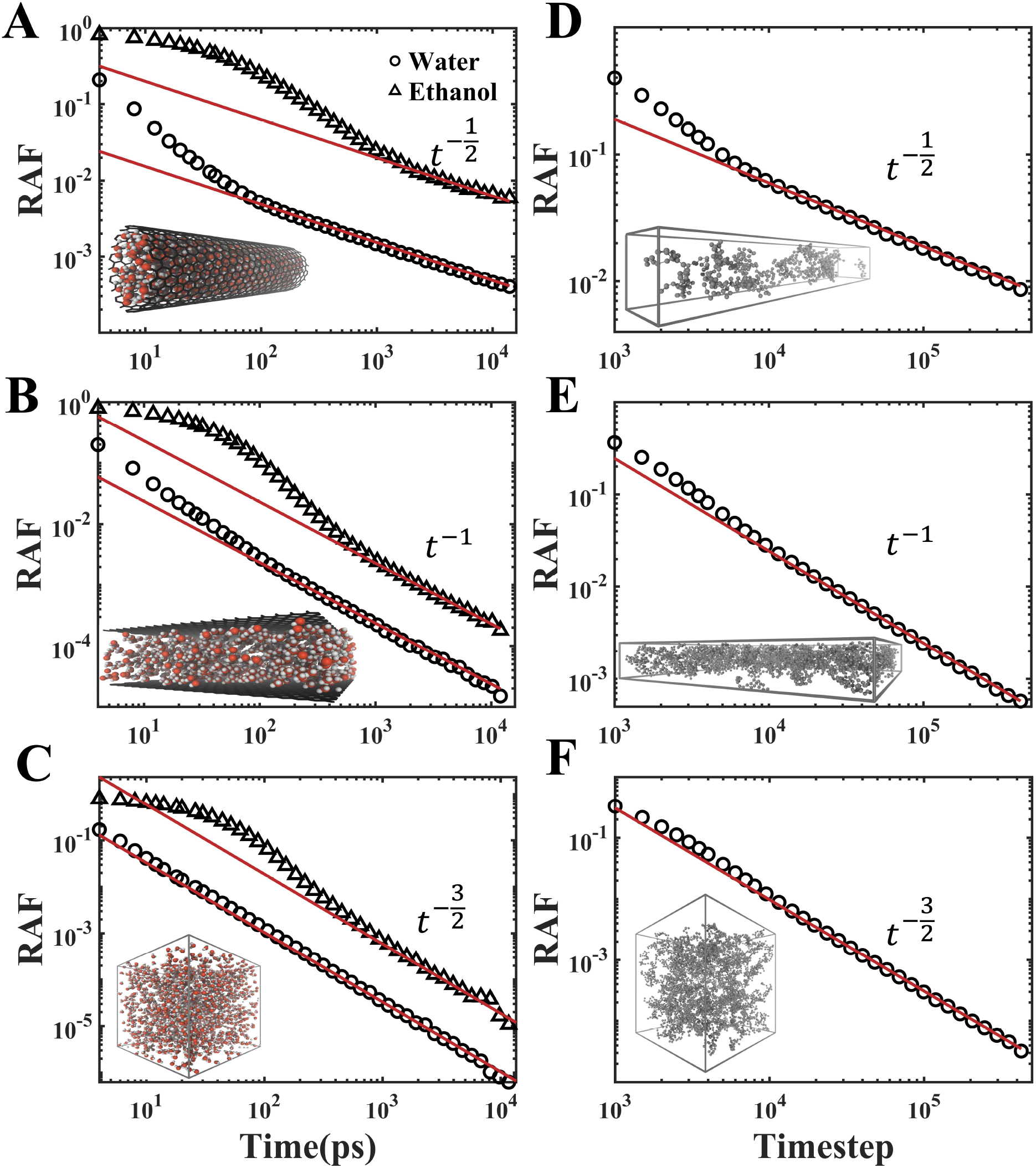
Universal scaling law of RAF in simple liquid. (**A-C**) Hydrogen bonding RAFs of water and ethanol in systems of different dimensions simulated by all-atom molecular dynamics. (**D-F**) RAFs of coarse-grained molecules in systems of different dimensions simulated by Brownian dynamics. The universal scaling law of *t*^-*D*/2^ is shown in the red lines as guide to the eye, where *D* is the dimension of the system. Typical snapshots of the systems are show in the insets of the panels.

**Fig. 2.**
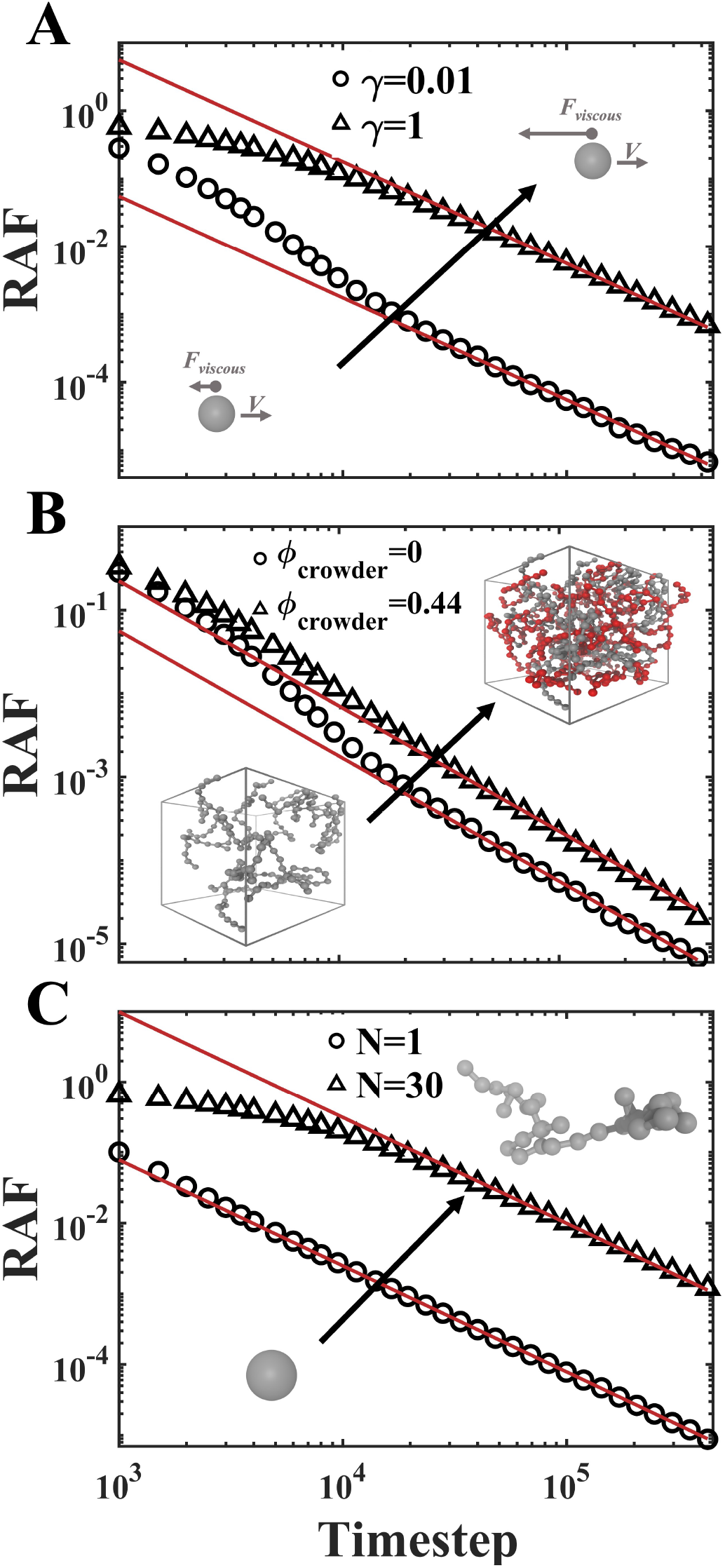
Factors that can augment the power law of RAF. The amplitude of the power law can be enhanced by increasing the viscosity of the liquid (**A**), the crowding level of the system (**B**), and the weight of the molecule (**C**). The universal scaling law is shown in the red lines as guide to the eye. The insets are schematic presentations of the changes of the systems.

In the continuum of affinities model, the power law of RAF emerges as the superposition of a wide spectrum of exponential RAFs with diverse binding affinities^15^. By tuning the shape of the spectrum, arbitrary power law scaling exponent can be achieved in theory. However, such tuning is not applicable if the RAF constituents are power laws sharing the same universal scaling exponent determined by the dimensionality of the system. The preservation of scaling under superposition is mathematically straightforward and verified by our simulations (Fig. 3). In this light, what surprises us from the TF experiments is not the reported RAF following a power law per se, but its scaling exponent being around -0.75, rending a much slower decay than the universal *t*^-1.5^ scaling expected in 3D space. Such discrepancy suggest that the anomalous RAF observed in TF experiments is nontrivial and cannot be resolved in the context of simple liquid. Understanding this anomaly might help us dissect the molecular mechanism and environmental complexity of trans-acting factors in gene regulation.

**Fig. 3.**
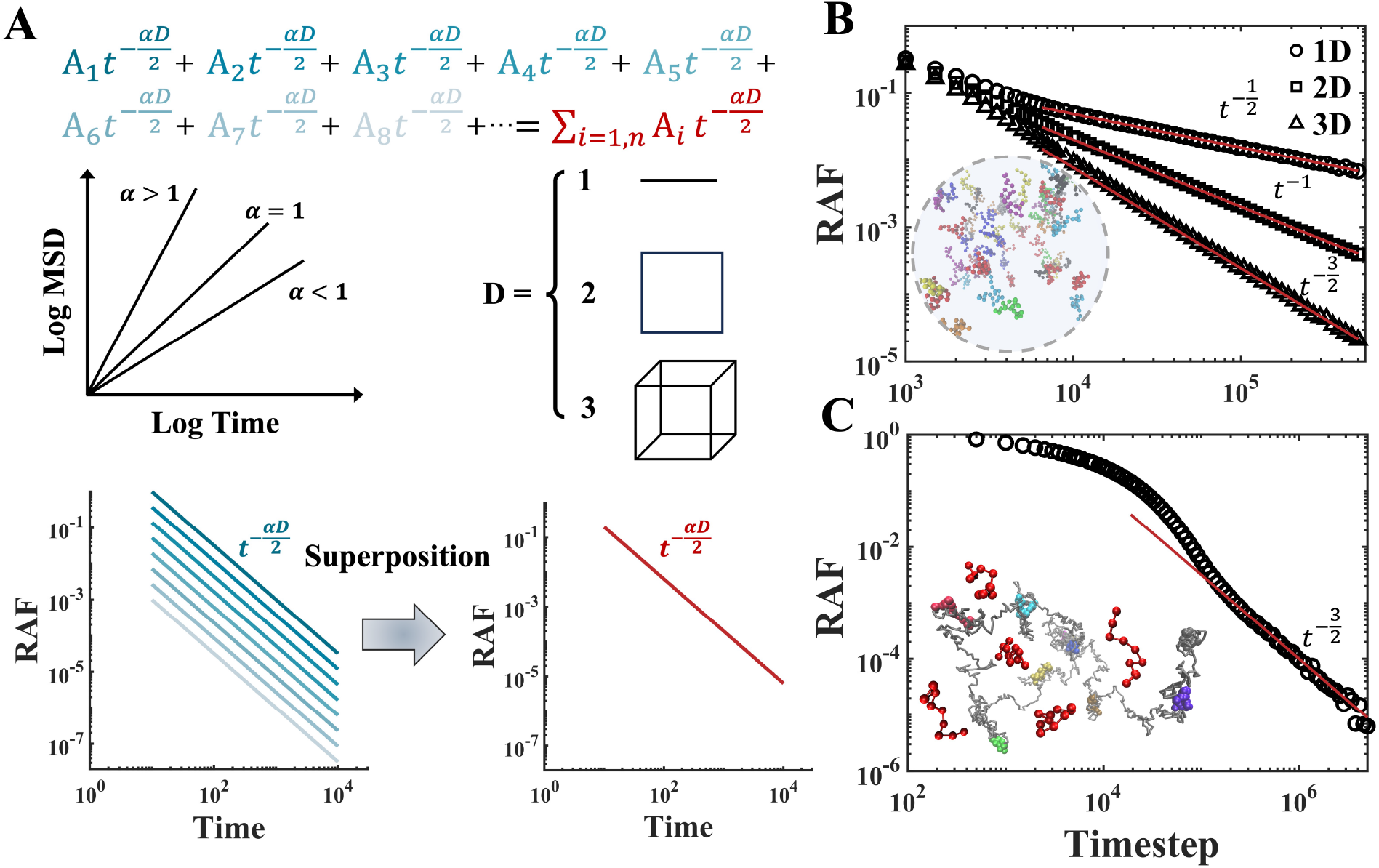
Preservation of the universal scaling law against affinity heterogeneity. (**A**) The superposition of power laws with the same scaling exponent does not change the scaling property of the result. In the equation, different amplitudes of the power law correspond to different affinities. In the scaling exponent, *α* stands for the diffusive exponent, and *D* is the dimension of the environment. See the Scaling theory section for the derivation of the power law. (**B**) Simulation verifies that diversifying the molecular types and the inter-molecular affinities has no effect on the scaling exponent of the RAF. The averaged RAF still follows the universal scaling law. Different molecules are marked by different colors in the inset. (**C**) Averaged RAF of trans-acting factors (red) binding to different CREs (other colors) of varying binding affinities on a dynamic chromatin polymer in a 3D liquid system. Again, simulation confirms that the asymptotic behavior obeys the universal scaling law.

### Scaling theory

Before we dive into more complex liquid environment, it is instructive to have some analytical insights for the power law scaling of RAF in simple liquid. While the exact RAF is difficult to analytically predict, it does not require too much mathematical work to obtain the scaling exponent of the asymptotic tail of the RAF. The simplest way to calculate this exponent may be through the scaling theory^34^, a powerful tool widely used in polymer physics.

It is clear that the asymptotic behavior of the RAF is determined by diffusion, and the effect of binding kinetics can be absorbed into a time-independent term at long-time scale that only contributes to the amplitude of the power law. This separation allows us to study the scaling exponent in the extreme case of vanishing binding energy. In this case the trajectory of liquid molecule can be mapped onto the conformation of a homopolymer. Therefore, analysis of the RAF is converted to solving the contact probability function *c*(*l*) that describes the likelihood of contact between two loci of the trajectory polymer as a function of the contour length between them. Here, the contour length *l* is equivalent to the time t in RAF.

The scaling property is associated with the concept of fractal^35^, i.e., self-similarity across scales. Not to loss generality, we assume that the trajectory polymer is of fractal dimension *θ* (the dimension of the walk), embedded in a *D*-dimensional space. The contact probability *c*(*l*) follows a power law of *l*^*s*^, where *s* is also the scaling exponent of RAF that we are interested in. Two loci can be said to be in contact if they are within a contact distance *b*, or *b*/*ε* in the measure of the scale unit *ε*. Changing the contact criterion will rescale the contact probability proportional to (*b*/*ε*)^*D*^. The contour length of a fractal is scale-dependent and can be written as *l*(*ε*) = *l*_0_*ε*^-*θ*^, with *l*_0_ being scale-invariant. Therefore, we have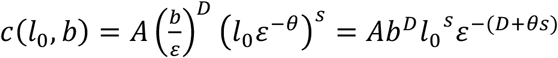, where *A* is a constant. Since *c*(*l*_0_, *b*) is scale-invariant, *D* + *θs* must vanish, leading to *s =* −*D*/*θ*. For normal diffusion we have *θ =* 2. In the case of anomalous diffusion, we have *θ =* 2/*α*, where *α* is the diffusive exponent. We refer to *s =* −*D*/2 as normal scaling, and in a more general situation that considers anomalous diffusion we have:

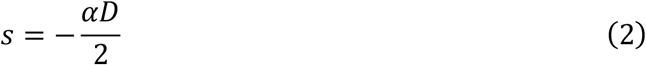

In simple liquid, the normal scaling of RAF (*s =* −1.5 in 3D space) is an asymptotic behavior that could be subtle and easily overlooked if the residence is highly transient^32,36^. In biological liquid, our simulations have indicated that the power law can be more prominent due to numerous factors (Fig. 2). Moreover, anomalous diffusion in biological systems^37^ can lead to anomalous scaling behavior of RAF. Typical proteins are subdiffusive^38,39^ with diffusive exponent around 0.8, which can lead to an anomalous RAF scaling of *t*^-1.2^. Such level of subdiffusion is, however, not sufficient to explain the *t*^-0.75^ scaling of TFs observed in experiment^12,15^. For random walk (*θ =* 2) in a fractal environment, one can prove based on the same scaling arguments that at long-time scale *s =* − *v*/2, where *v* is the fractal dimension of the environment. From an anomalous diffusion point of view (Eq. 2), we have *α* = *v*/*D*. Given the fact that the environmental fractal dimension is around 2.5 for the cell nucleus^40,41^, the corresponding scaling exponent of -1.25 is still inconsistent with the TF experiment. The discrepancy suggests that there are other mechanisms at play behind the strong anomaly of TF residence.

### Effects of aggregation and phase separation

So far, we have focused on simple liquid that is single-phase and passive. Biological system, in contrast, is multi-phase in nature and active in chemical reactions. We reason that, it is this complexity of the liquid environment rather than the heterogeneity of binding energies that gives rise to the anomalous RAFs of the TFs observed in experiments. To interrogate this possibility, we study RAF in phase-separated liquid. We use small droplets as the condense phase because nuclear hubs related to transcription are often of limited sizes^42–44^. In fact, whether the formation of small hubs in the cell nucleus is driven by LLPS, the mechanism that unmixes oil and water, has been debated intensively^44–46^. To avoid any bias, we also investigate the RAF of a non-LLPS hub, which we refer to as aggregate in oppose to the droplet in the LLPS case. In our simulation, both the aggregate and the droplet are recruited to a CRE on a segment of chromatin polymer. One essential difference between the two is that, after the recruitment, turning off the affinity of the CRE will dissolve the aggregate but not the droplet (Fig. S9).

In our simulation of the aggregate (Fig. 4A), we find that the RAF does not break the universal *t*^-1.5^ scaling law (Fig. 4B). However, when LLPS happens (Fig. 4C), the decay of the RAF becomes remarkably slower. A *t*^-1.06^scaling appears and dominates a considerable duration of time before it is succeeded by a faster decay (Fig. 4D). To understand the effect of droplet on the molecular residence time, we decompose the RAF into two parts:

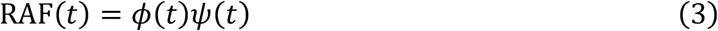

**Fig. 4.**
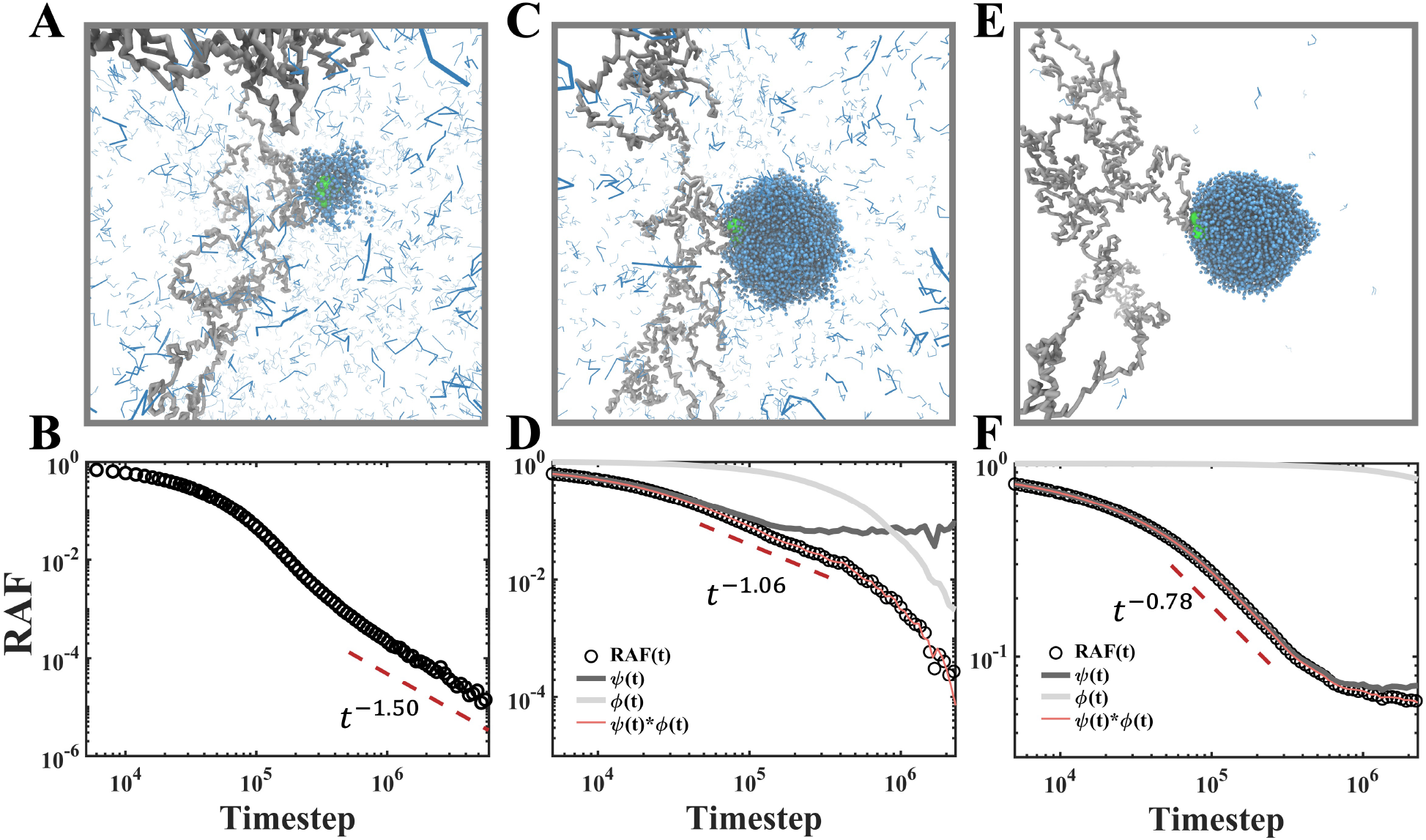
Breakdown of the universal scaling law in passive LLPS systems. (**A, B**) Snapshot and RAF signature in an aggregate system without LLPS. The CRE is marked in green and the trans-acting factors in blue. The aggregate of the trans-acting factors is unstable without the CRE. (**C, D**) Snapshot and RAF signature in a weak LLPS system with low partition coefficient between the two phases. (**E, F**) Snapshot and RAF signature in a strong LLPS system with high partition coefficient between the two phases. Decomposition of the RAFs according to Eq. 3 are shown in the LLPS systems. Dashed lines of power laws are added as guide to the eye.

For a molecule that binds to the CRE at *t* = 0, *ϕ*(*t*) characterizes the probability of finding this molecule inside the droplet as a function of time. The faster the droplet exchanges mass with its exterior, the faster *ϕ*(*t*) decays with time. In experiment, the mass exchange property of the droplet is usually measured by the FRAP technique^47^. The second function *ψ*(*t*) describes the time-dependent probability of this molecule residing near the CRE, given the condition that it stays inside the droplet at this moment. The decomposition shows that the droplet has a caging effect on the molecules within it, which slows down the decay of RAF. However, such caging is not absolute since the droplet quickly exchanges its mass with the environment, leading to a fast decay of RAF at longer time scale. Remarkably, we find that the *t*^-1.06^scaling of the RAF is not sensitive to the binding energy between the molecules and the CRE, if the degree of phase separation is fixed (Fig. S11). In order to reproduce the anomalously slow *t*^-0.75^ scaling in the TF experiments, we strengthen the inter-molecular affinity to intensify the phase separation (Fig. 4E). At the price of a very high partition coefficient (the concentration ratio between dense and dilute phases), we are able to obtain a scaling of *t*^-0.78^as shown in Fig. 4F. However, the time window of this scaling is less than one order of magnitude, much shorter than the experimental observation that is almost two orders of magnitudes^12,15^. At larger time scale, the simulation reports a flatter decay of the RAF, due to the repressed mass exchange between the dense phase and the dilute phase. At even larger time scale, we expect the RAF to decay faster as the mass exchange starts to kick in.

The above simulation results demonstrate that non-LLPS aggregation has limited effect on the scaling of RAF. In contrast, LLPS readily breaks the universal scaling law but still fails to explain the anomalous scaling behavior of TFs reported by experiment^12,15,16^. This discrepancy between simulation and experiment could have two opposite interpretations: (1) TFs do not undergo LLPS in vivo, or (2) endogenous TFs are subject to a new type of LLPS that differs from the unmixing of oil and water. We decide the second possibility is more intriguing and worth of further exploration.

### Emergent scaling in active droplet

Many nuclear proteins, including the TFs and RNA-binding proteins (RBPs), carry intrinsically disordered regions (IDRs) that are prone to phase separation and post-translational modifications (PTMs)^5,9,16,48–54^. According to in vitro experiments, some IDRs and IDR-containing proteins can undergo liquid-to-gel transition^55–57^, suggesting that the weak inter-IDR affinity could strengthen over time. Such passive aging effect, in combination with the enzyme-catalyzed reaction of PTM, is expected to further complicate the multi-phase biological liquid and reshape the RAF of molecular factors in living cells. To study this problem, we build an active droplet in-silico by adding both positive and negative feedback loops to the phase separation mechanism (Fig. 5A). The positive feedback is included implicitly to describe the aging of inter-molecular affinity, whereas the negative feedback is modeled explicitly by adding enzyme molecules to the system (see the Methods for more information). The condensation of the substrate molecules will recruit the enzyme molecules to the surface of the droplet. A snapshot of the active droplet is presented in Fig. 5B, with distinct molecular states marked by different colors. The proportions of different molecular states are shown in the color bar. It is worth noting that the system is out of thermodynamic equilibrium, yet in a steady state. For simplicity, all the molecules in the droplet have the same binding affinity to the CRE of the chromatin, regardless of their states. We analyze the molecular residence near the CRE of the chromatin and plot the RAF in Fig. 5C.

**Fig. 5.**
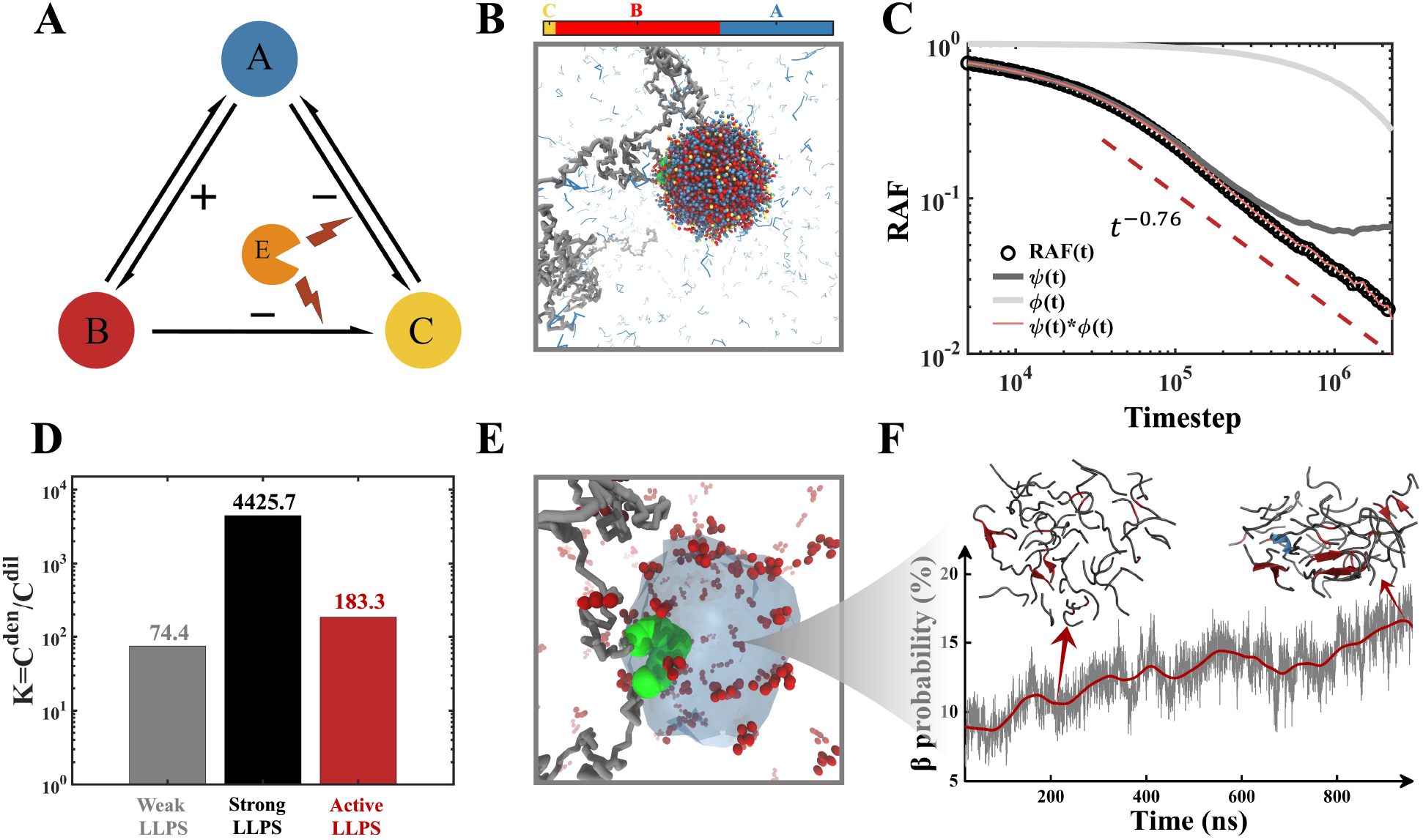
Emergence of anomalous scaling in an active droplet. (**A**) Design principles of the feedback loops in the dissipative LLPS system. The three states from A to C correspond to normal trans-acting factors, aged factors with increased inter-molecular affinities, and post-translationally modified factors with lower inter-molecular affinities, respectively. Positive and negative signs represent the counteracting feedback loops. The enzymes are only implicated in the negative feedback loop. (**B**) Simulation snapshot of the active droplet. The color bar represents the proportion of the three molecular states. (**C**) The RAF and its decomposition according to Eq. 3. The anomalous scaling in line with TF experiment is demonstrated by the dashed red line as guide to the eye. (**D**) Partition coefficients between dense and dilute phases in different LLPS systems shown in log-scale. The active LLPS maintains a low partition coefficient. (**E**) Typical simulation snapshot shows that the enzymes (marked in red) for the negative feedback control tend to reside at surface of the active droplet. (**F**) Zoomed-in view of the droplet aging process that serves as a positive feedback loop. Atomistic simulation shows that, in a model system of 40 human hnRNPA1 segments (residues 243-248), the density of *β*-structure grows with time, which increases the inter-molecular affinity and further condenses the droplet. The *β* -structure is highlighted in red in the snapshots.

To our surprise, a prominent power law scaling appears, spanning over almost two orders of magnitudes in time (Fig. 5C). More interestingly, the scaling exponent of the power law happens to be -0.76, in good agreement with TF experiments^12,15^. Note in experiments, different TFs exhibit nonidentical RAFs, but most of their scaling exponents nonspecifically lie within -0.75±0.10, which robustness is hard to comprehend in the continuum of affinities model^15^. To make sure that the emergent scaling phenomenon in our simulation is also robust, we carry out an independent in-silico experiment with modified affinity and reaction parameters in our model. We find that, despite the redistribution of different molecular states in the active droplet, the scaling exponent of RAF is not sensitive to the change of the system (Fig. S13). The robust anomality of the RAF scaling is remarkable as it only emerges in nonequilibrium phase-separating systems regulated by counteracting feedback loops.

Compared to passive phase separation without feedback, the active droplet manages to reconcile the conflict between long residence lifetime and fast mass exchange with the dilute phase (Fig. 5C) that maintains low partition coefficient between phases (Fig. 5D). This is achieved by balancing the aging and the metabolic PTM reactions of the phase-separating molecules. While the PTM reactions tend to occur at the surface of the droplet where the enzymes are enriched (Fig. 5E), affinity aging can happen anywhere of the droplet. This aging process could be driven by the growing formation of protein secondary structure such as *β*-sheet^58^, that is implicated in the liquid-solid phase transition of some nuclear proteins. Using all-atom molecular dynamics simulation,we show that the density of *β*-sheets grows as the concentration of peptides increases (Fig. 5F), confirming that secondary structuring can provide a positive feedback loop in LLPS.

### Scaling proofreading model

The emergent scaling law of RAF in active droplet is a fascinating phenomenon in physics, nevertheless, biologists might be more interested in its potential function in living cells. Here, we propose a scaling proofreading mechanism to elucidate the functional advantage of the anomalous RAF scaling of TFs (Fig. 6). Just like high specificity is required in biosynthetic processes^59,60^, selective residence of TFs at target DNA sequences is critical for gene regulation. In the intricate landscape of the eukaryotic genome, characterized by a multitude of non-specific binding sites, mere passive diffusion and binding prove insufficient to confer the high specificity necessary^3,61^. To address this conundrum, we posit that the long-time tail of the RAF is functionally more important than the average residence time of the TFs. In other words, it is the small TF population of long residence time and higher rebinding probability that plays decisive role in gene regulation. Compared to the exponential decay of RAF assumed in canonical models, power law decay revealed by our simulation has a more pronounced long-time tail. Among power laws of different scaling exponents, the differentiation between their survival probabilities will be augmented with time, allowing higher selectivity at long-time scale. However, such augmentation mechanism is impossible in simple liquid, where the scaling exponent of RAF is fixed by the dimensionality of the system. In a sense, such universal scaling at equilibrium condition is analogous to a ground state. Upon active phase separation that costs energy, the scaling exponent of RAF gets excited to a larger value, which significantly increases the population of TFs of long residence times. Taking advantage of such non-continuous jump in the scaling exponent, the transcription machinery can readily filter out the non-specific binding events whose RAF scaling exponents stay at the ground state, despite of their varying binding affinities. Consuming energy to increase the TF specificity, the active LLPS therefore effectively function as a proofreading mechanism to reduce the error of gene regulation. The scaling proofreading model suggests that by integrating amplification and dissipation into LLPS, evolution can harness the thermodynamic force unmixing oil and water to shape the flow of genomic information.

**Fig. 6.**
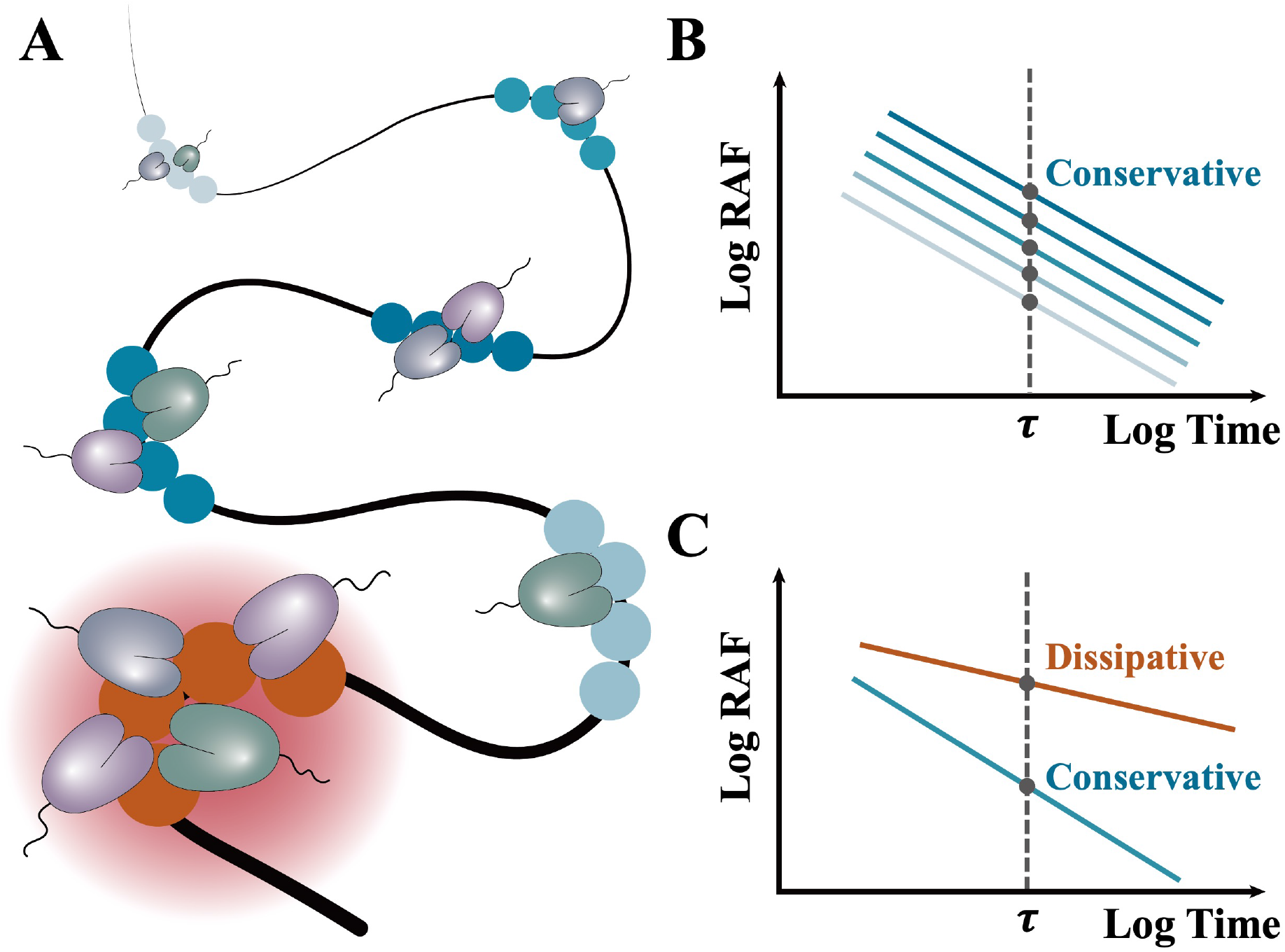
Schematics of the scaling proofreading mechanism. (**A**) TFs bind to genomic loci of varying binding affinities on the chromatin polymer. The binding loci without LLPS are marked in blue with the darker colors corresponding to stronger affinities. The binding loci with active LLPS is shown in red. (**B**) In the absence of LLPS, the RAFs from all the loci at long-time scale obey the universal scaling law, regardless of the binding affinity. Therefore, the specificity is poor and does not improve over time. (**C**) Active LLPS consumes energy to create an anomalous RAF scaling that is much slower than the universal scaling law. Over the time, the difference between the two scaling functions becomes more pronounced, allowing for high specificity of TFs in gene regulation. In this scaling proofreading process, the specificity is provided by active LLPS rather than strong binding affinity.

## Discussion and conclusions

The trajectory of a liquid molecule over time is like a polymer conformation in space, whose statistical properties encode rich environmental information. The exciting experimental progress in molecular imaging and tracking has made it possible to precisely measure the RAF of nuclear proteins^7,11^. Deciphering such experimental data provides a non-invasive means to probe the complex nuclear environment, but prior knowledge is needed. By systematically simulating a variety of liquid systems, we have shown that the fundamental assumption of exponential RAF in biophysics is invalid. Power law scaling of RAF dominates single-phase liquid (Fig. 1-3), breaks down in passive multi-phase liquid (Fig. 4) and reemerges in active multi-phase liquid (Fig. 5).

Normal phase separation at thermodynamic equilibrium tends to be macroscopic, but only microscopic droplet can lead to strong caging effect for the RAF. In our simulations, the microscopic size of the droplets is realized by controlling the total number of proteins in the simulation box. It merits a note that, the active droplet can constrain its own size due to the negative feedback loop that is absent in the passive droplets.

We should stress that the scaling laws in passive and active liquid are of different nature. The former is a universal law in a conservative system, whereas the latter is an emergent phenomenon in a dissipative system^62–65^. Remarkably, the two systems exhibit drastically different scaling exponents, and it is the dissipative system that reproduces the anomalous scaling exponent of TFs observed in experiments. These results imply that endogenous LLPS is not only an important driving force to condensate nuclear proteins, but also a heavily regulated process subject to feedback loops.

Compared to negative feedback mechanisms that repress LLPS^54,66,67^, positive feedback loops are largely overlooked. We show that affinity aging can serve as a positive feedback mechanism to stabilize the residence of functional molecular factors in the active droplet (Fig. 5). Our finding indicates that the driving force behind the phase transition from a liquid to an amyloid state^68^, commonly associated with neurodegenerative diseases^27,69,70^, may possess intrinsic physiological functions in gene regulation. Such duality underscores the complexity of the biological phase transition and prompts further exploration into the intricate feedback mechanisms that govern its role in both pathological and normal cellular contexts.

Our simulation results lead to the hypothesis of a scaling proofreading mechanism of TF selectivity, which links residence scaling anomaly to functional advantage. In contrast to the continuum of affinities model, the scaling proofreading model suggests that the power law of RAF is not a result of genome-wide superposition at thermodynamic equilibrium, but an all-or-none phenomenon emerges locally from nonequilibrium phase separation. Such sensitive phase behavior provides a biophysical basis for high TF fidelity in gene regulation against the noise from non-specific binding. Our model predicts that the residence scaling properties of TFs can be modified by changing their affinities to their cognate condensates, which can be tested by experiment in future.

There are a few limitations of our model. For simplicity, we have only considered one kind of component (with three states in the case of active droplet) in our phase separating system as a proof of concept. In living cells, biological phase separation is heterogeneous in nature, involving multiple components. In our coarse-grained model, we have not accounted for the effects of physical interactions on the local chemical equilibria of the reactions. These factors and more molecular details are worth further investigation but should not change the general conclusions in this work.

In summary, the intricate interplay between kinetics, dynamics, and phases can lead to rich, non-exponential RAF behaviors of liquid molecules. Power laws exist in both conservative and dissipative systems but with distinct scaling exponents. Our understanding of the emergent scaling anomaly of TFs unfolds a living picture of biological condensates that acquire function through metabolism and aging.

## Methods

Analysis of the RAF is a computationally expensive task for large-scale simulations. Employing a hash table for the neighbor list along with OpenMP parallelization, we boost the speed of RAF analysis by nearly three orders of magnitudes. Amending the artifact of periodic boundary condition (PBC) will inevitably lead to loss of samples over time. To resolve this conundrum, we introduce image binding sites to ensure the correct asymptotic behavior and high statistical quality of the RAF.

We use all-atom molecular dynamics simulations conducted in GROMACS to study the RAFs of water and ethanol. For one-dimensional confinement, molecules are restricted within a single-walled carbon nanotube of 2.1 nm in diameter, with PBC applided to the axial direction. In the case of two-dimensional confinement, we use two parallel graphene sheets separated by 2.0 nm, applying PBC to the directions parallel to the graphene surfaces. The bulk system is simulated in a cubic box of 16 nm with PBC in all directions.

In our simulation of the affinity aging mechanism, all proteins are modeled using the AMBER99SB-ILDN force field, and water molecules are represented by the TIP3P model. Initially, 40 peptides (GYNGFG) are randomly distributed in a cubic box of 10 nm size. After energy minimization and equilibration, 1 microsecond of simulation with the NPT ensemble is conducted for data production.

Brownian dynamics (BD) using LAMMPS is carried out to study the RAFs of coarse-grained proteins in larger systems. In the simulation of single-phase liquid, the coarse-grained proteins consist of 20 beads interconnected by harmonic springs. The non-bonded isotropic interactions between all monomers are modeled using the standard 12-6 Lennard-Jones potential. In the simulation of multiphase liquid, the chromatin segment is simplified as a polymer of 1500 beads including a 20-bead binding site, and proteins are represented as 5-bead chains.

In our model of active phase separation, the protein monomer can be in one of three states: A, B or C, corresponding to the normal state, the state with aged or amplified affinity, and the state with repressed affinity after PTM, respectively. The probability of transition from A to B increases with the local protein concentration, leading to a positive feedback. Such transition is reversible. The transition from A to C is a PTM reaction catalyzed by enzyme molecules that are explicitly modeled in our system. The rate of this reaction is designed to be proportional to the local concentration of the enzyme. The transition from C back to A is modeled implicitly and predominantly occurs in the dilute phase. Both positive and negative feedback loops of LLPS are implemented into an in-house version of LAMMPS.

## Acknowledgements

We acknowledge financial support from the Shenzhen Bay Laboratory, the Shenzhen Bay Laboratory Open Fund Project (SZBL2021080601013 to K.H.), and computational support from the Shenzhen Bay Laboratory Supercomputing Center. We thank Ming Yang, Lin Li and Xiangli Liu for assisting the project.

## Author contributions

Conceptualization: K.H. Theory and model: K.H. Methodology: S.Q., Z.Y., and K.H. Investigation: S.Q., Z.Y., and K.H. Analysis: S.Q. and Z.Y. Funding acquisition: K.H. Project administration: K.H. Supervision: K.H. Writing – original draft: K.H. Writing – review and editing: S.Q., Z.Y., and K.H.

