## Supporting Information for "Scaling laws of molecular residence time"

The PDF file includes:

Supplementary Text

Figures. S1 to S14

### A. Single-phase system

#### All-atom liquid system

We constructed different liquid environment for water and ethanol, applying different dimension of confinement. For one-dimensional (1D) confinement, a single-walled carbon nanotube with $2.1\,nm$ in diameter was employed to restrict the movement of liquid molecules, incorporating periodic boundary conditions along the z-axis. In the case of two-dimensional (2D) confinement, the setup involved two infinite parallel graphene sheets separated by a distance $L_z = 2.0\,nm$, with periodic boundary conditions implemented in both y and z directions. In the bulk water system, periodic boundary conditions were uniformly implemented across all directions. The integration time step was 2 fs and coordinates were saved every 4 ps. During the relaxation before data production, temperature is kept at 298.15K and it is controlled with Nose-Hoover thermostat. Our all-atom molecular dynamics (MD) simulations were conducted with GROMACS software package[1], utilizing the Lennard-Jones parameters from the OPLSAA force field in GROMACS for modeling the interactions between carbon, water, and ethanol.

#### Liquid system without hydrodynamics

The flow and movement of surrounding fluid could either facilitate or impede the approach and retreat of molecules, thereby affecting the dynamics of intermolecular bonding. To validate that the power-law tail observed in the long-term behavior of hydrogen bonds is not a result of hydrodynamic factor, we designed a simulation system based on Langevin dynamics, where hydrodynamics effects are not considered. In this simulation, molecules were coarse-grained into polymers, each consisting of N beads interconnected by harmonic springs, defined by $U_b(\mathbf{r}) = k(r - r_0)^2$.In this context, $k$ represents the rigidity of the connected spring and is assigned a value of 70, while $r_0$, the equilibrium bond distance, was fixed at $1\sigma$. The non-bonded isotropic interactions between all monomers were modeled using the standard 12-6 Lennard-Jones (LJ) potential, $U_{LJ}(r) = 4\epsilon\left[\left(\frac{\sigma}{r}\right)^{12} - \left(\frac{\sigma}{r}\right)^{6}\right]$, to accurately represent molecular interactions. Different values of $N$ were employed to examine molecules of various molecular weights, and the Lennard-Jones interaction parameter $\epsilon$ between monomers was also varied to investigate binding affinities of different strengths. Additionally, a virtual reflective wall was introduced in a specific dimension to selectively confine particle movement along that axis. For all coarse-grained simulation systems, the initial configuration first undergoes $2 \times 10^5$ simulation steps equilibration with all interactions turned off. After then, the interactions are turned on and a relation of $1 \times 10^6$ simulation steps is performed. Another period of $1 \times 10^6$ simulation steps is used for data production. Trajectories are saved each 500 steps for subsequent analysis.

### B. Multi-phase system

**Passive phase separation**

To explore our hypothesis that the experimentally observed power-law residence time behavior stems from the phase separation occurring at chromatin binding sites, we adopted a bead-spring model for chromatin representation and simulated protein motion and interaction with chromatin. Specifically, the chromatin fiber is modeled as a polymer chain with 1500 coarse-grained beads, featuring a binding site composed of a sequence of 20 consecutive beads. Proteins are depicted as chains of 5 beads, each connected sequentially by bonds. In this simulation, when no specific interaction occurs between the binding site and the protein, the protein tends to distribute evenly across the entire simulation box. Conversely, if there is an inducing interaction between the binding site and proteins, a heterogeneous distribution occurs, which can be further classified into two types. In the first case, at protein concentrations below saturation, the binding site, through a strong specific interaction, induces a small-sized aggregate. In the second scenario, at higher concentrations, beyond the saturation point but potentially still in a metastable state of phase separation, the inducing effect of the binding site enables the system to overcome the nucleation energy barrier, facilitating phase separation. This differs from the previously mentioned aggregate in that, when the interaction between the binding site and the protein is turned off, the aggregate dissipates, whereas the phase-separated droplets remain stable, albeit detaching from the binding site location (see Figure S9).

**Active phase separation**

In the simulation of active regulation of phase separation, the monomers in the protein polymer can be in one of three states-A, B or C. ‘A’ monomers correspond to the normal state, state ‘B’ to the enhanced interaction state, frequently triggered by intermolecular secondary structures, and state ‘C’ represents a reduced interaction state, commonly, as result of protein modifications like phosphorylation or glycosylation. The incorporation of a positive feedback chemical reaction $A \leftrightharpoons B$ is achieved by dynamically changing the monomer type between ‘A’ and ‘B’. The switch rate is not constant but instead depends on the surrounding concentration of protein monomers. Specifically, the probability of forward reaction is positively correlated with the nearby protein concentration, reflecting the increased propensity for forming intermolecular secondary monomers at higher concentrations. Conversely, the shift from B to A is less likely to occur as the concentration increases. Central to our model is a regulating enzyme that can be recruited to the surface of condensates, thereby selectively altering the intermolecular interactions of proteins at this interface. The implementation of this modification of affinity in our simulation is achieved through a probabilistic switching of protein monomers from state A to C. The rate of this reaction is designed to be directly proportional to the local concentration of the enzyme. The reversal of post-translational modifications, specifically from state C back to A, which we implicitly simulate, predominantly occurs outside the droplets. To integrate both positive and negative feedback loops into the phase separation model, we have developed an in-house version of Large-scale Atomic/Molecular Massively Parallel Simulator (LAMMPS)[2].

**C. Residence time analysis**

To characterize the survival time of intermolecular bonding, we define a residence time autocorrelation function $\mathrm{RAF}(t)$, which is $\mathrm{RAF}(t) = \frac{\langle n(t)n(0)\rangle}{\langle n(0)n(0)\rangle}$. In this expression, $n(t)$ represents the bond formation status, having a value of 1 when a pair is formed between two molecules, and zero otherwise. The determination of hydrogen bond formation is based on geometric criteria, where a bond is considered formed when the distance between two molecules is less than 3.5Å and, simultaneously, the donor-hydrogen-acceptor angle $\theta_{DHA}$ is greater than 130 degrees. All the hydrogen bond lifetime analysis in our work were performed using the Python library MDAnalysis[3]. To analyze the lifetime of intermolecular binding within the framework of Langevin dynamics, we developed our own analysis code, underpinned by a distance-based criterion. Given the large number of bonded pairs in complex systems, such as in three-dimensional and phase-separated environments, we optimized several key aspects to accelerate out data processing. To address the need for constructing an intermolecular contact matrix at every time interval T, we created keys by reading the neighbor list of each molecule. This approach enabled the use of a hash table to efficiently identify the active units in the contact map at each time point, thereby providing complete matrix information. Secondly, to enhance the efficiency of our data analysis, we parallelized our code using OpenMP. Furthermore, in our study of chromatin-TF binding dynamics, we faced a significant data-quality issue related to the rapid migration of proteins out of the simulation area. This phenomenon caused a continuous decline in the sample size over time, as slower-moving binding sites remained within the simulation box. This effect, combined with the inherently limited availability of binding sites, led to increased data noise. To address this issue, we introduced image binding sites in our analysis. This approach effectively counteracted the loss of proteins from the simulation domain, significantly improving the reliability and robustness of our results.

In our analysis of RAF, another noteworthy consideration is the appropriate selection of coordinates when determining the binding status between two molecules under periodic boundary conditions. Commonly in molecular dynamics simulations, wrapped coordinates are used for constructing neighbor lists. However, for accurate binding status assessment in our study, it is essential to use unwrapped coordinates instead. The use of wrapped coordinates in this context can lead to a misinterpretation of the periodic boundaries as fixed, resulting in incorrect dynamics interpretation (see Figure S2). All our self-developed analysis techniques and the customized LAMMPS code are available on GitHub (https://github.com/zhijs/autocorrelation-for-lammps).

**D. Theory**

In the main text we used a scaling theory to obtain the general relationship between the RAF scaling exponent and the properties of the system (Eq. 2). If we do not consider anomalous diffusion and fractal dimension, this relationship is simply s=-D/2, which can be proved using other mathematical techniques. Consider a particle undergoing one-dimensional Brownian motion with position $x(t)$. The movement of the particle is characterized by a diffusion coefficient D. The probability distribution $P(x,t)$ can then be described by the diffusion equation, also known as the Fokker-Planck equation:

$$\frac{\partial P(x,t)}{\partial t} = D\frac{\partial^2 P(x,t)}{\partial x^2} \qquad (S1)$$

Assume that the particle starts as the origin, $x = 0$, at time $t = 0$. Then the corresponding initial condition can be written as:

$$P(x,0) = \delta(x) \qquad (S2)$$

The diffusion equation can be solved with this condition. To do this, we apply the Fourier transform on both sides of the eq. (1) with respect to x:

$$\frac{\partial \mathcal{F}\{P(x,t)\}}{\partial t} = D\mathcal{F}\left\{\frac{\partial^2 P(x,t)}{\partial x^2}\right\} \qquad (S3)$$

$$\frac{\partial F(k,t)}{\partial t} = -Dk^2F(k,t) \qquad (S4)$$

where $F(k,t) = \int P(x,t)\exp(-ikx)\,dx$. Now we obtain a first-order differential equation in time for $F(k,t)$. To solve this, we use the method of separation of variables:

$$\frac{\partial F(k,t)}{F(k,t)} = -Dk^2\partial t \qquad (S5)$$

Integrating on both sides and we get:

$$lnF(k,t) = -Dk^2t + C(k) \qquad (S6)$$

where $C(k)$ is a constant. And we can get the expression for $P(k,t)$: $P(k,t) = e^{-Dk^2t+C(k)}$. According to initial condition in real space (Eq. S2), we can determine the initial condition in the Fourier space by taking the Fourier transform of $\delta(x)$:

$$F(k,0) = \int \delta(x)\exp(-ikx)\,dx = \exp(-ik\cdot 0) = 1 \qquad (S7)$$

Now we can obtain the value of $C(k)$ and write down the solution of $F(k,t)$:

$$F(k,t) = \exp(-Dk^2t) \qquad (S8)$$

To find $P(x,t)$, we need to take the inverse Fourier transform of $F(k,t)$:

$$P(x,t) = \frac{1}{2\pi}\int \exp(-Dk^2t)\exp(ikx)\,dk = \frac{1}{\sqrt{4\pi Dt}}e^{-\frac{x^2}{4Dt}} \qquad (S9)$$

We then expand this probability density function of the displacement to a 3D random walk. In each direction, the particle follows a separate independent random walk. In 3D, the probability density function $P(\mathbf{r},t)$ can be expressed as a product of three 1D probability density functions, one for each dimension:

$$P(\mathbf{r},t) = P(x,t)\cdot P(y,t)\cdot P(z,t) = \frac{1}{\sqrt{4\pi Dt}}e^{-\frac{x^2}{4Dt}}\cdot\frac{1}{\sqrt{4\pi Dt}}e^{-\frac{y^2}{4Dt}}\cdot\frac{1}{\sqrt{4\pi Dt}}e^{-\frac{z^2}{4Dt}}$$

$$= \frac{1}{(4\pi Dt)^{\frac{3}{2}}}e^{-\frac{r^2}{4Dt}} \qquad (S10)$$

Now we try to relate the Brownian motion of particles to the dynamics of bonds between particles. Assuming that both particles undergo one-dimensional Brownian motion, the probability of their relative position falling within the interaction range can be regarded as the probability of bond formation, To explore this, we first consider the probability of each individual particle being at a specific location:

$$P_1(x_1,t) = \frac{1}{\sqrt{4\pi Dt}} e^{-\frac{x_1{}^2}{4Dt}} \quad (S11)$$

$$P_2(x_2,t) = \frac{1}{\sqrt{4\pi Dt}} e^{-\frac{x_2{}^2}{4Dt}} \quad (S12)$$

The relative distance between these two particles is $x_{rel} = x_2(t) - x_1(t)$, and the probability distribution of the relative position, $P_{rel}(x_{rel}, t)$ can be expressed as:

$$P_{rel}(x_{rel},t) = \int_{-\infty}^{+\infty} P_1(x_1,t) \cdot P_2(x_1 + x_{rel}, t) dx_1 \quad (S13)$$

Substitute the expression for $P_1(x_1,t)$ and $P_2(x_2 + x_{rel}, t)$ and gets:

$$P_{rel}(x_{rel},t) = \frac{1}{4\pi Dt} \int e^{-\frac{x_1{}^2}{4Dt}} e^{-\frac{(x_1+x_{rel})^2}{4Dt}} dx_1 \quad (S14)$$

We can rewrite the integrand in $P_{rel}(x_{rel},t)$:

$$P_{rel}(x_{rel},t) = \frac{1}{4\pi Dt} \cdot \left[-\exp\left(\frac{x_{rel}{}^2}{8Dt}\right)\right] \int_{-\infty}^{+\infty} e^{-\frac{\left(x_1+\frac{1}{2}x_{rel}\right)^2}{2Dt}} dx_1 \quad (S15)$$

Using the Gaussian integral property, we can directly write down the fina result:

$$P_{rel}(x_{rel},t) = \frac{1}{\sqrt{8\pi Dt}} \cdot \left[-exp\left(\frac{x_{rel}{}^2}{8Dt}\right)\right] \quad (S16)$$

Notice that Eq. S15 is based on the assumption that both two particles starts at the zero point. A more general expression for $P_{rel}(x_{rel},t)$ accounting for nonzero initial positions $x_1(0)$ and $x_2(0)$, can be formulated as:

$$P_{rel}(x_{rel},t) = \frac{1}{\sqrt{8\pi Dt}} \cdot \left[-exp\left(\frac{x_{rel}{}^2 - \left(x_1(0) - x_2(0)\right)^2}{8Dt}\right)\right] \quad (S17)$$

Since our study focuses on the survival time of these intermolecular bonds, we choose to observe particles that initially maintain a bond (i.e., the initial bond status is intact). This implies that the relative distance between these particle pairs is initially within the scope of the interaction range. Then we can define the probability of bond reformation:

$$P_{reform} = \int_0^{x_c} P_{rel}(x,t)dx = \int_0^{x_c} \frac{1}{\sqrt{8\pi Dt}} \cdot \left[-exp\left(\frac{x^2}{8Dt}\right)\right] dx \quad (S18)$$

where $x_c$ is the interaction radius. When the interaction range is considerably shorter compared to the scale of particle motion (i.e., long $t$ and small $x_c$), the exponential term in Eq. S17 approaches 1, yielding the final expression for 1D confinement:

$$P_{reform}^{1D} = \frac{1}{\sqrt{8\pi Dt}} \quad (S19)$$

Likewise, the probability of bond reformation for 2D and 3D cases can be written as:

$$P_{reform}^{2D} = (8\pi Dt)^{-1} \quad (S20)$$

$$P_{reform}^{3D} = (8\pi Dt)^{-\frac{3}{2}} \quad (S21)$$

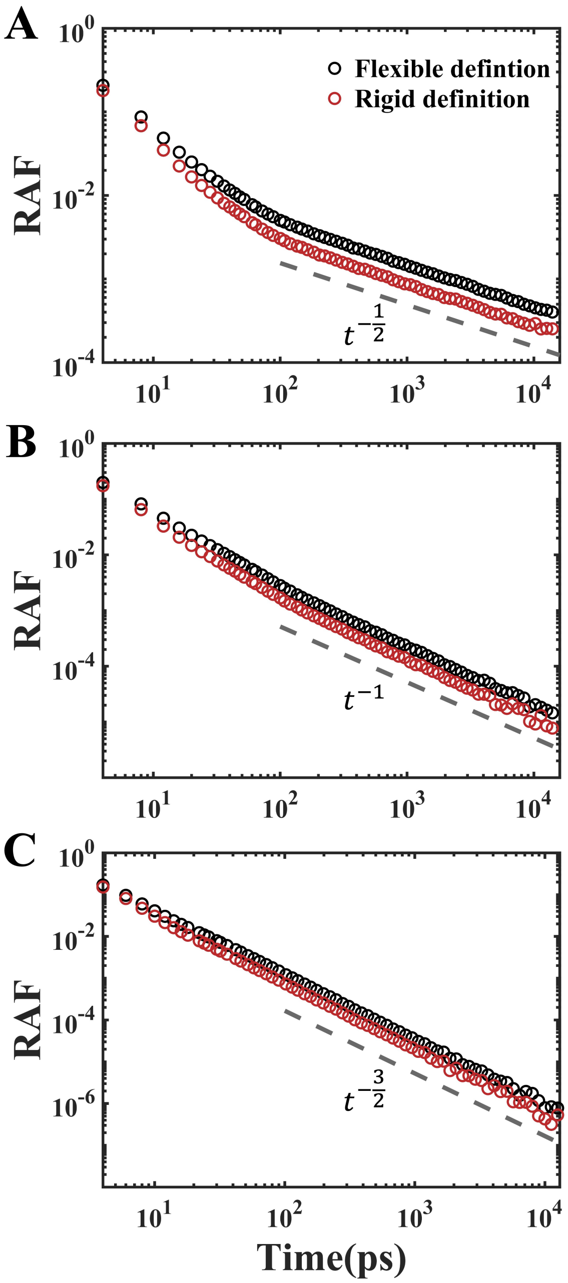


Figure S1. Residence time auto correlation function RAF of hydrogen bonding for different water confinement: (A) 1D-confined, (B) 2D-confined, and (C) bulk water. Two hydrogen bond criteria are applied in calculating RAF: a conventional criterion (bonding if intermolecular distance < 3.5Å and donor-hydrogen-acceptor angle > 130°), represented by black lines, and a stricter criterion (distance < 3.0Å and $\theta_{DHA} > 150°$), shown by red lines. Dashed gray lines in each plot are used to guide the eye for the asymptotic behavior of $t^{-\frac{1}{2}}$, $t^{-1}$, and $t^{-\frac{3}{2}}$.

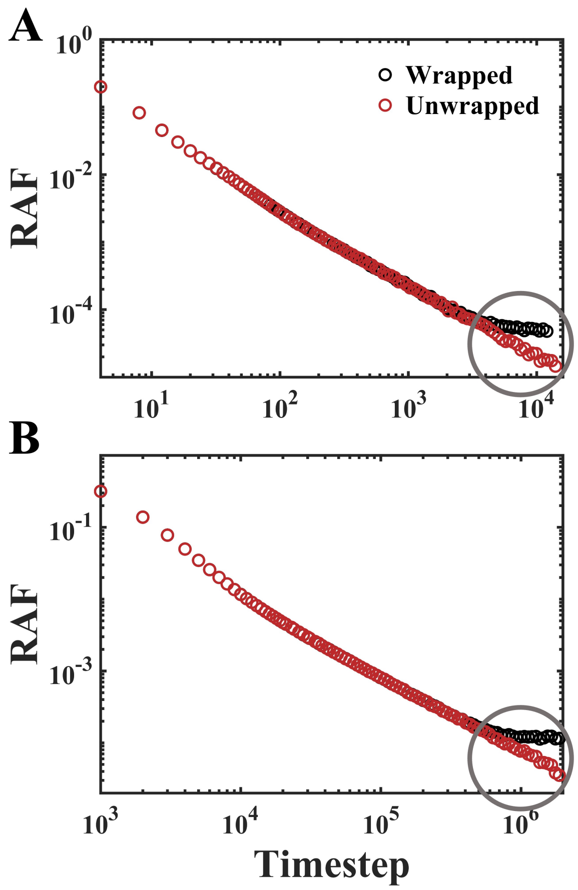


Figure S2. Analysis of RAF by using wrapped (black) and unwrapped (red) molecular coordinates in (A) all-atom and (B) coarse-grained homogeneous phase systems. The key differences are highlighted on the long-time scale, marked by grey circles. At these stages, molecules reach the periodic boundaries. The application of wrapped coordinates in this context effectively converts the originally periodic directions into fixed boundaries, which can lead to wrong dynamics interpretation.

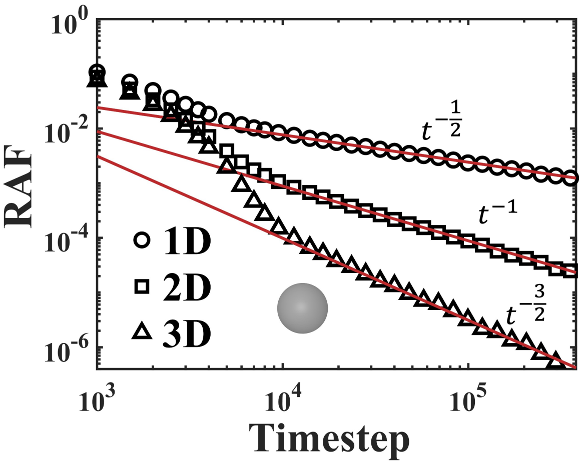


Figure S3. RAFs in systems of different dimensions, simulated by Langevin dynamics. This system is modeled as a single-bead model, where each bead specifically represents an individual molecule.

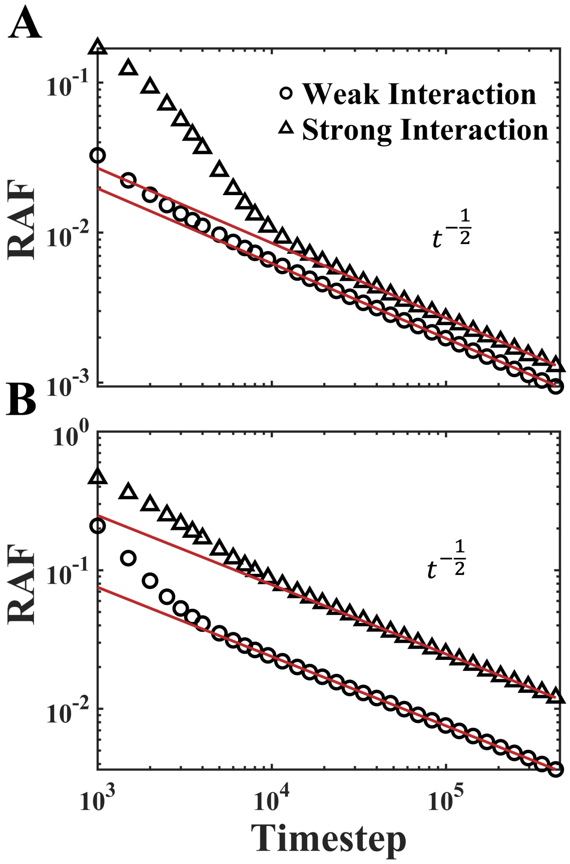


Figure S4. Log-log plot of the autocorrelation function RAF in (A) a single-bead model, and (B) a bead-spring polymer model based on Langevin dynamics. These simulations were conducted in a 1D-confined system. The graph demonstrates lines with points of varying shapes, indicative of different binding strengths (with $\epsilon$ changing from 0.05 to 0.50). A stronger molecular affinity results in a higher intensity of RAF and delays the onset of the power-law region, although the overall asymptotic power-law behavior remains unchanged.

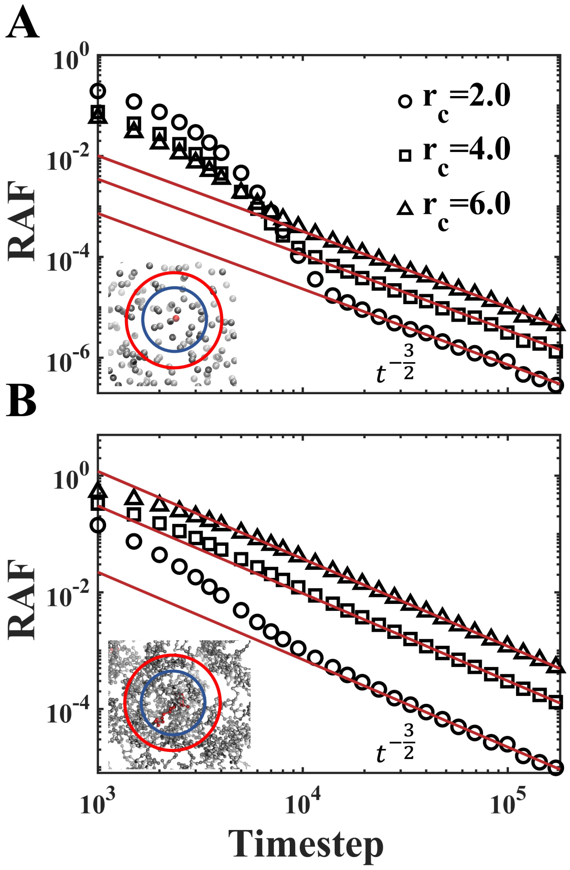


Figure S5. Log-log representation of the autocorrelation function RAF in (A) a single-bead model, and (B) a bead-spring polymer model based on Langevin dynamics. These in-silico experiments were conducted in a 3D bulk system. The lines with different shaped points in the graph represent the use of varying cutoff distances in defining bond formation. In the inset, we present schematic representations of different residence range selections. The choice of different residence ranges affects the intensity of RAF but does not modify its asymptotic power-law behavior.

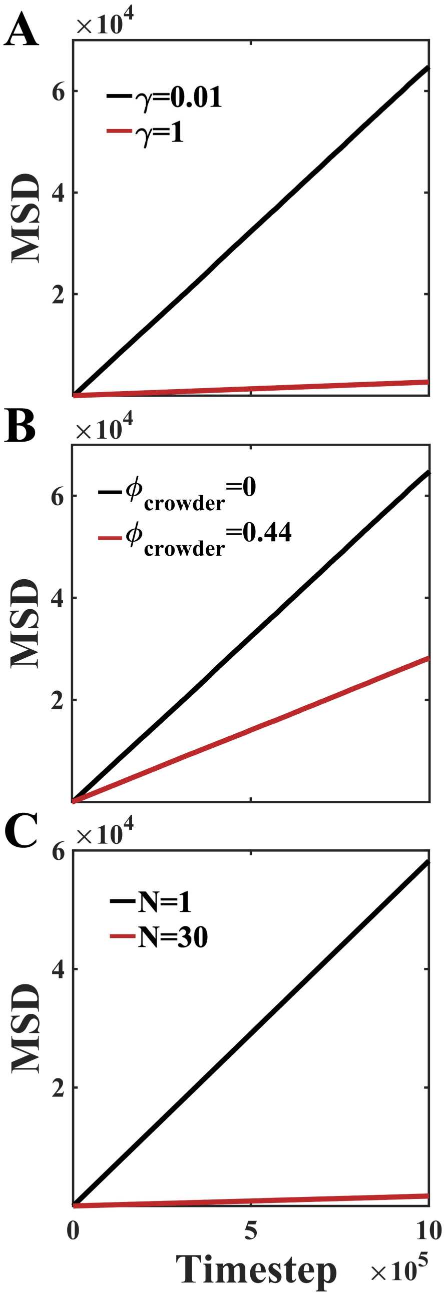


Figure S6. Temporal evolution of the mean-squared displacements (MSD) of molecules in environments with different viscosity, crowding, and molecular weight conditions. These factors significantly influence molecular diffusion, which in turn affects the autocorrelation function of bond formation.

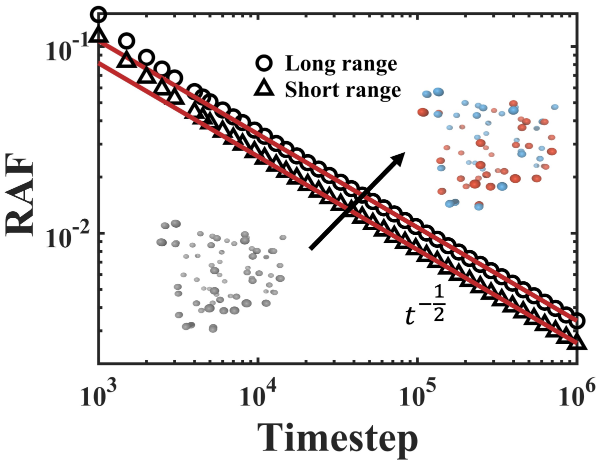


Figure S7. Time evolution of the bond autocorrelation function RAF in the presence of long-range electrostatic forces. In this computational experiment, a 1D-confined system was chosen to optimize computational efficiency. Additionally, the original single-bead model is divided into an equal number of positively and negatively charged beads. It is observed that short- or long-range nature of intermolecular forces does not affect the power-law behavior at long timescales.

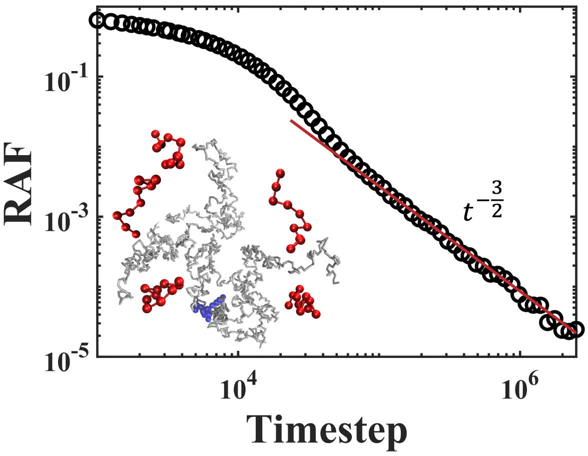


Figure S8. Autocorrelation function of transcription factors' residence time at specific chromatin binding sites. Unlike in Figure 3(B), this experiment investigates the scenario where the transcription factor has a single type of binding affinity with specific binding sites. The inset provides a schematic of the in-silico experimental system. Solid red lines labelled $t^{-\frac{3}{2}}$ are added to guide the eye.

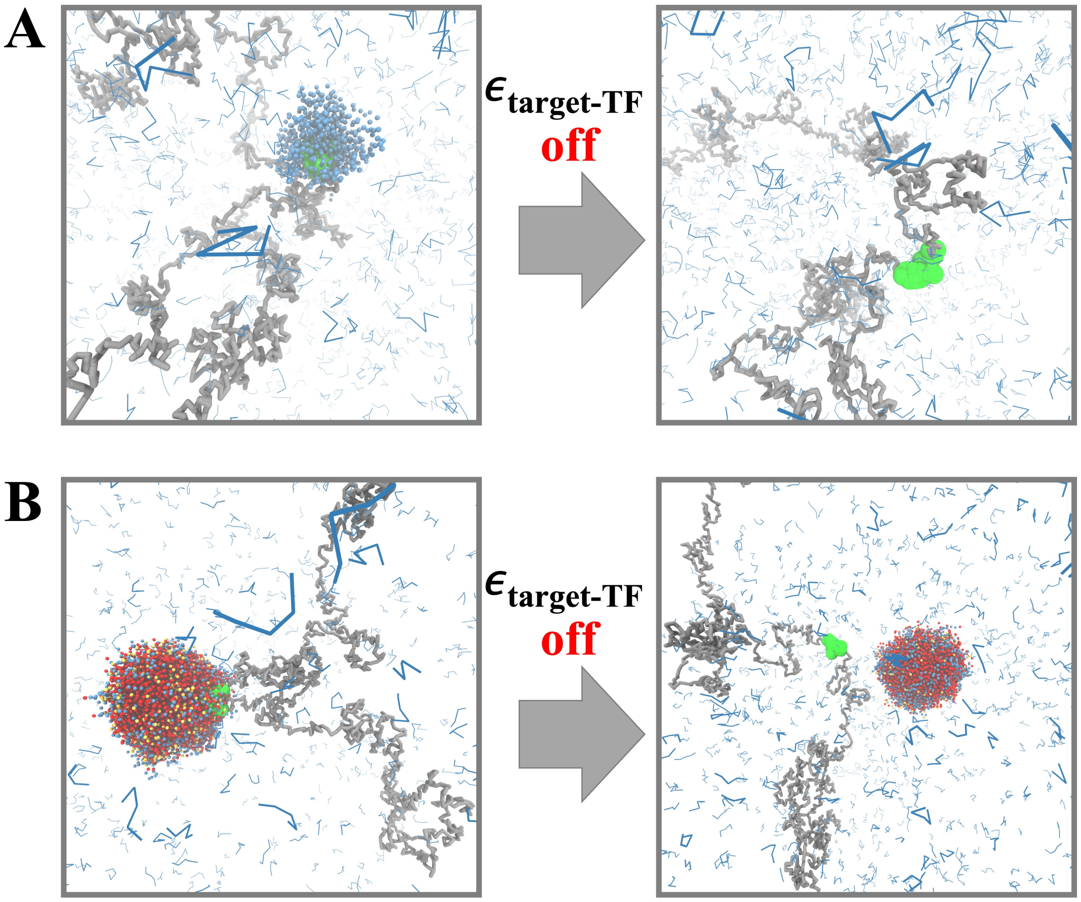


Figure S9. (A) Simulation configuration of the aggregate system before and after deactivating CRE-TF interactions. (B) Configurational changes in the droplet system upon turning off CRE-TF interactions. These observations highlight significant differences between the two systems when CRE-TF specific interactions are turned off. In the aggregate system, the condensate dissipates upon deactivation of the interactions. However, in the droplet case, the droplet remains intact but becomes detached from the CRE.

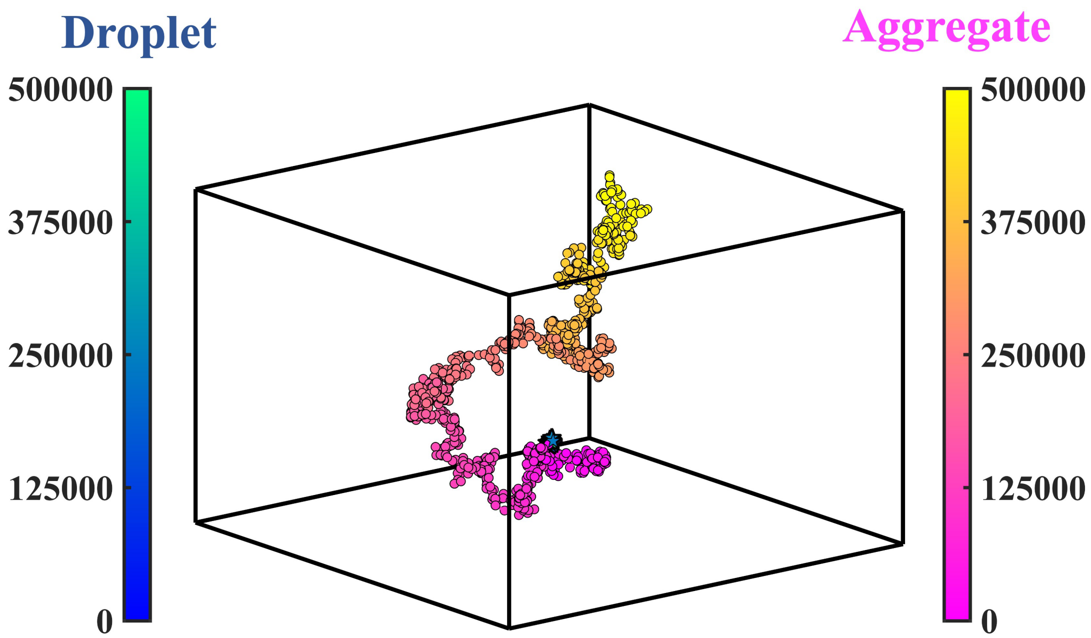


Figure S10. In both aggregate and droplet systems, we selected typical trajectories of transcription factors that resides at the CRE at t=0 and depicted them in two distinct colors in this figure. The ‘winter color’ represents single-molecule trajectories in the droplet scenario, while the ‘spring color’ indicates those in the aggregate system. These two systems exhibit significant differences. During an observation period of 500000 time units, the droplet system shows limited large-scale displacement as molecules are confined within droplets. In contrast, in the aggregate system, transcription factors, after residing in small-sized condensates for a short-time period, start a random walk.

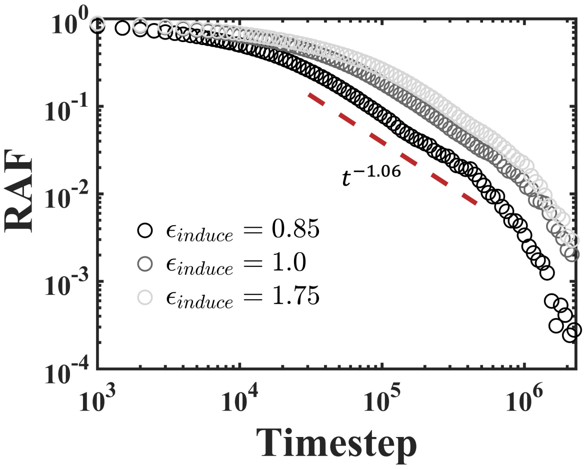


Figure S11. RAF in a passive phase-separated system with a constant interaction strength of $\epsilon = 0.53$ between TFs, but different CRE-TF affinities. The interaction strengths between CRE and TF are set at $\epsilon = 0.85, 1.0$ and 1.75. Higher CRE-TF affinity lead to a slower decay in the short-term dynamics, predominantly governed by an exponential function, leading to a generally stronger RAF. However, the long-term power-law behavior and the exponents remain unchanged.

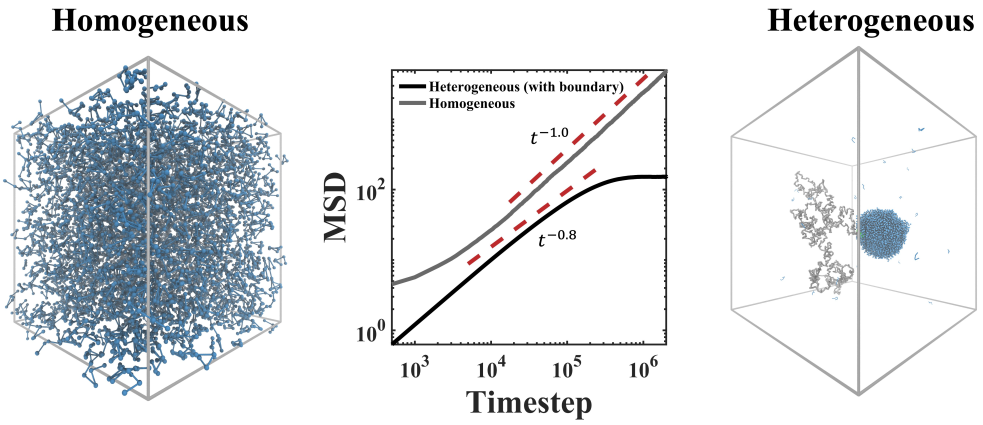


Figure S12. Characterization of TF diffusive behavior in homogeneous, self-crowded environments, and heterogeneous phase-separated systems, as determined by the mean-squared displacement over time in a log-log scale. In the self-crowded system, proteins initially exhibit a subdiffusive regime (MSD~$t^{\alpha}$ with $\alpha < 1$), which later crosses over to a normal diffusion (MSD~$t$). TFs in phase-separated systems predominantly exhibit subdiffusion over a prolonged intermediated phase, eventually reaching a plateau in MSD, indicative of confinement effects at the droplet interface.

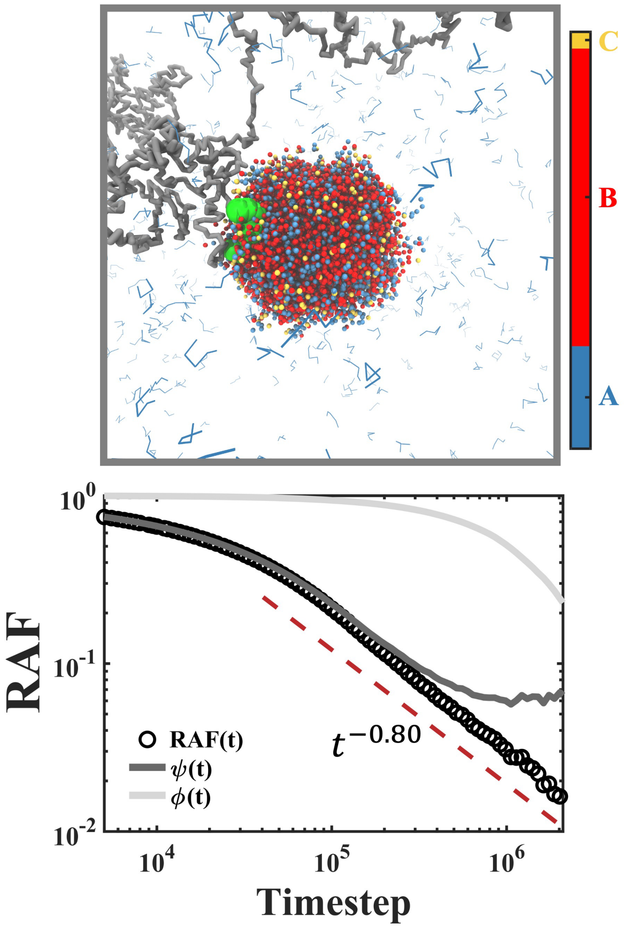


Figure S13. Simulation snapshot in an active LLPS system, with different molecular states illustrated by different colors. The proportion of these states are displayed in the color bar. Differing from Figure 5 (B), this simulation applies another set of parameters for reaction rate and protein interaction strengths. The RAF is characterized and decomposed. Dashed red line labelled $t^{-0.80}$ is added to guide the eye.

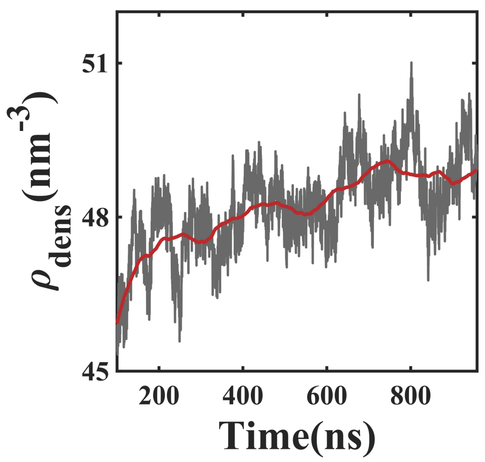


Figure S14. Temporal density evolution of the dense phase during aggregation process of 40 human hnRNPA1 residues (243-248), predicted by all-atom MD simulation.
